# Novel insights into the economic relevant phenotypes of *Bombyx mori* strains from Romanian breeding centres

**DOI:** 10.1101/2024.06.14.598941

**Authors:** Gabriela-Maria Baci, Adrian Ionascu, Attila Cristian Ratiu, Daniel Severus Dezmirean

## Abstract

The silkworm, *Bombyx mori*, has an inestimable economic and pharmaceutical value, but it is also a very effective model organism for biological and biotechnological sciences. Herein, we characterize several phenotypic parameters particular for Băneasa 1 (B1) and Galben de Băneasa (GB) Romanian silkworm breeds, as well as of Auriu Chinez (ACH) and JH3 breeds originating from China and, respectively, Japan. Our analysis revealed concrete differences and similarities between the phenotypic properties of the four breeds, allowing their grouping consecutive to PCA and KODAMA analyses. Our study is the first to characterize the four breeds regarding their biological competencies and offers a glimpse at various ventures aiming at the employment and amelioration of these silkworm breeds.

## 1. Introduction

*Bombyx mori* holds a prominent position in the global economy by exerting its impact across various fields, including agriculture, pharmaceuticals, and biomedicine. Arguably the greatest sericultural by-product that exhibits a tremendous impact on the field of life sciences is the fibroin protein. Fibroin is a key polymer found in the silk thread produced by the silkworm.

The silk fibroin structure comprises of a light chain (Fib-l) and a heavy chain (FIBH) associated at the C-terminus through a single disulfide bond. A glycoprotein (P25) is linked to this complex through noncovalent interactions (Inoue et al., 2000). The assembled ratio of the three components is 6:6:1, which forms a structure of approximately 2.3 MDa. Silk fibroin is synthesized in the posterior silk gland (PSG) of the last larval instar, following transfer to the middle part of the silk gland (MSG). Whilst residing in MSG, the molecules of sericin are added and the resulting complex is transported to the anterior part (ASG) to form the silk thread (Inoue et al., 2000).

The Fib-l and FIBH polypeptides are encoded by *Fib-l* and *FIBH* genes localized on chromosome 14 and 25, respectively (Gamo, 1982). *FIBH* has one intron and two large exons, while *Fib-l* gene has seven exons separated by large introns. The latter gene mostly contains non-coding DNA as the first intron represents 60% of the gene, and the rest of the introns add up to 31% (Barbosa et al., 2009).

Currently, for using silk fibroin as a biomaterial, investigating the mechanisms of fibroin production are of significant research interest, in order to influence its properties or to widen its spectrum of applicability. For example, Xu et al. (2018) replaced the *FIBH* gene with the *major ampullate spidroin-1* gene from the spider *Nephila clavipes*, thus considerably enhancing the extensibility of the silk threads produced by the silkworms (Xu et al., 2018). Table 1 summarizes various studies that obtained silk fibers with superior properties by genetically engineering the silkworms.

**Table 1.**
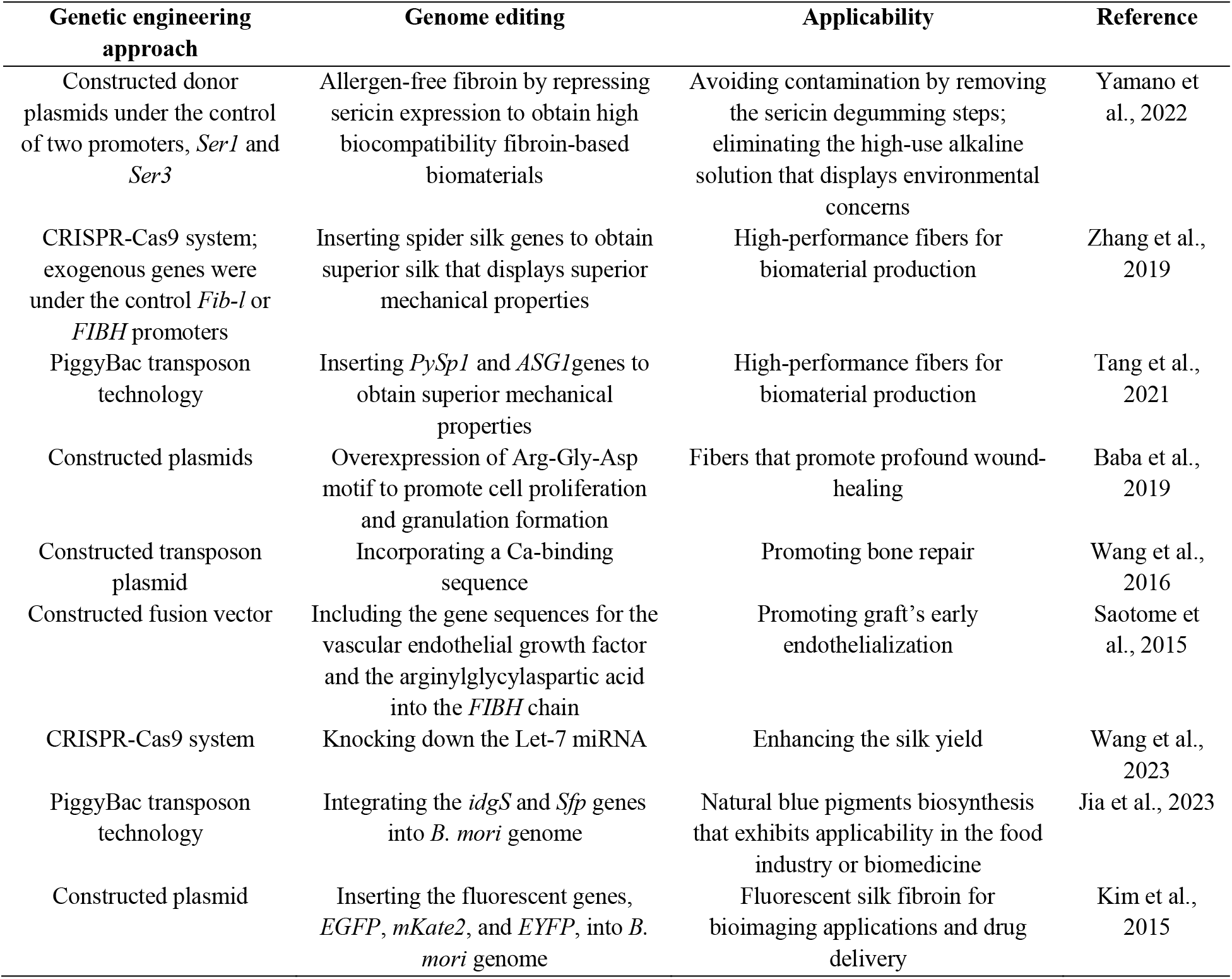
Summarization of various genome editing approaches leading to specific applicability

In addition to the importance of its sericultural by-products, the silkworm attracted a lot of interest from researchers due to its great potential for producing recombinant proteins. By knocking out the endogenous *Fib-l* or *FIBH* gene, an important increase in terms of recombinant protein production has been reported (Tatematsu et al., 2010).

Herein, we characterize various biological and economic parameters of Băneasa 1 (B1) and Galben de Băneasa (GB) Romanian silkworm breeds, as well as of Auriu Chinez (ACH) and JH3 breeds originating from China and, respectively, Japan. Our investigation revealed a comprehensive image of the four *B. mori* breeds’ potential economic performances.

## 2. Materials and Methods

### 2.1 Breeds maintenance

Four silkworm breeds (*B. mori*) were evaluated in this study, specifically B1 and GB, two Romanian breeds, JH3 Japanese breed, and ACH, a Chinese breed. The breeds that exhibit foreign origins, specifically, JH3 and ACH, were reared under the climatic conditions of Romania. Disease-free eggs were obtained from the Global Centre of Excellence for Advanced Research in Sericulture and Promotion of Silk Production (International Sericultural Commission structure) from the University of Agricultural Sciences and Veterinary Medicine of Cluj-Napoca, Romania. Eggs incubation was performed under hygienic conditions at 24 ± 1°C and 75–80% humidity. The experiment intended to evaluate various traits of larvae instars, was carried out in November, and due to the lack of fresh mulberry leaves, the silkworms were reared on artificial diet obtained from the Agricultural Academy Scientific Centre on Sericulture (Vratsa, Bulgaria). A standard strategy was adopted for silkworm rearing, specifically in the first instar, as the larvae were reared at 28°C with 80-90% humidity. The temperature was decreased by 1°C with each instar development, and, in the last two larval instars, the humidity was reduced to 70-80%.

### 2.2 Assessment of Biological and Economical Parameters

Several larvae instar parameters were evaluated, including the egg incubation period, larva length and weight, as well as the larval duration. Furthermore, on the 5^th^ day of the 5^th^ instar, the weight of the silk glands was assessed. After the fulfillment of the cocoon spinning period, a series of traits were measured, namely the incubation period (days), larval weight (g), larval instars duration (days), silk gland weight (g), larval length (mm), cocoon transverse axis length (mm), fecundity (No.), pupae weight (g), cocoon longitudinal axis length (mm), and cocoon weight (g). For each target parameter, the measurements were performed on ten individuals from each breed, that were randomly chosen. Raw data of measured biological parameters is provided in the Supplementary Table.

### 2.3. Data analysis

Statistical analysis and graphical representations of assessed biological parameters data were performed using the R programming language version 4.3.1 (R Core Team, 2023) and the RStudio IDE version 2023.06.2-561 (RStudio Team, 2020). An original R script was designed for data analysis (data not published) based on the *dplyr* (Wickham *et al*., 2023), *ggplot2* (Wickham, 2016), *nortest* (Gross and Ligges, 2015), *tseries* (Trapletti and Hornik, 2023), *car* (Fox and Weisberg, 2019), *stats* (R Core Team, 2023), *FactoMineR* (Le *et al*., 2008), *factoextra* (Kassambara and Mundt, 2020) and *KODAMA* (Cacciatore et al., 2014) packages. Statistical significance was considered for p < 0.05 at 95% confidence.

## 3. Results

Prior to performing statistical testing on the biological parameters measured for the individuals pertaining to the analyzed *B. mori* breeds, we investigated the summary statistics and normality of the data. The mean and standard error (SE) of each variable are presented in Table 2.

**Table 2.** Mean and its standard error (SE) for the measured parameters for the considered *B. mori* breeds.

| Breed | B1 |  | JH3 |  | GB |  | ACH |  |
| --- | --- | --- | --- | --- | --- | --- | --- | --- |
|  | Mean | SE | Mean | SE | Mean | SE | Mean | SE |
| Incubation period (days) | 9.947 | 0.162 | 10.88 | 0.169 | 12.4 | 0.152 | 13.06 | 0.098 |
| Larval weight (g) | 3.019 | 0.025 | 2.724 | 0.023 | 2.305 | 0.026 | 2.252 | 0.016 |
| Larval duration (days) | 32.74 | 0.24 | 32.89 | 0.312 | 33.95 | 0.135 | 34.11 | 0.186 |
| Silk gland weight (g) | 0.8275 | 0.103 | 0.807 | 0.096 | 0.698 | 0.092 | 0.675 | 0.086 |
| Larval length (mm) | 51.45 | 0.202 | 50.38 | 0.25 | 49.8 | 0.292 | 48.76 | 0.373 |
| Cocoon transverse axis length (mm) | 15.2 | 0.157 | 15.37 | 0.151 | 15.18 | 0.172 | 14.59 | 0.256 |
| Fecundity (No.) | 390.5 | 11.729 | 387 | 15.497 | 318.1 | 10.179 | 401.1 | 15.359 |
| Pupae weight (g) | 0.389 | 0.021 | 0.352 | 0.018 | 0.361 | 0.023 | 0.325 | 0.016 |
| Cocoon longitudinal axis length (mm) | 32.23 | 0.319 | 31.11 | 0.377 | 30.26 | 0.763 | 30.52 | 0.362 |
| Cocoon weight (g) | 0.482 | 0.024 | 0.412 | 0.019 | 0.445 | 0.028 | 0.436 | 0.016 |

In order to adjust the statistical analyses based on data distributions, data normality was evaluated by performing a series of statistical tests, such as Kolmogorov–Smirnov and Shapiro–Wilk tests. Graphical representations in the form of histograms and quantile-quantile (Q-Q) plots were used in association with normality testing (Supplementary Figures 1 and 2). We identified that data corresponding to the majority of the measurements are normally distributed therefore we opted to perform parametric statistical tests.

We evaluated the differences between the four local breeds of *B. mori* by performing an analysis of variance on the independently measured parameters (Table 3). Regardless of the statistical test used in our analysis (the parametric one-way ANOVA test which assumes equal variances, the parametric Welch’s ANOVA test which takes unequal variances or the nonparametric Kruskal-Wallis test), we observed statistically significant differences between the breeds for the majority of measured parameters. The differences between breeds for “Cocoon longitudinal axis length” were considered statistically significant, showing a p value of 0.055 for the one-way ANOVA test, 0.006 for the Welch’s ANOVA test, and 0.019 for the Kruskal-Wallis test. Interestingly, the “Cocoon transverse axis length” was not significantly different between the four breeds for the one-way ANOVA test (p=0.15) but showed significance in the case of nonparametric testing (p=0.029). Given the nature of the data for this parameter, normal distribution, and unequal variances, we considered that Welch’s ANOVA results were most suitable in this case, resulting in no significant differences between the four breeds. Only three measured parameters did not display statistically meaningful distinctions between the *B. mori* breeds, namely “Cocoon transverse axis length”, “Pupae weight”, and “Cocoon weight”.

**Table 3.** Results of one-way ANOVA, Welch’s ANOVA and Kruskal-Wallis statistical testing between the four *B. mori* breeds for each measured parameter. Statistical tests were performed in R.

| Statistical test | One-way ANOVA |  | Welch's ANOVA |  | Kruskal-Wallis test |  |
| --- | --- | --- | --- | --- | --- | --- |
| Statistic value | F value | p value | F value | p value | $\chi^2$ | p value |
| Incubation period (days) | 88.58 | $< 2 \times 10^{-16}$ | 103.02 | $< 2.2 \times 10^{-16}$ | 55.411 | $5.612 \times 10^{-12}$ |
| Larval weight (g) | 11.75 | $2.93 \times 10^{-6}$ | 10.712 | $3.674 \times 10^{-5}$ | 25.279 | $1.35 \times 10^{-5}$ |
| Larval duration (days) | 8.512 | $7.37 \times 10^{-5}$ | 8.415 | $2.686 \times 10^{-4}$ | 21.312 | $9.069 \times 10^{-5}$ |
| Silk gland weight (g) | 7.458 | $2.25 \times 10^{-4}$ | 7.88 | $3.692 \times 10^{-4}$ | 19.177 | $2.5 \times 10^{-4}$ |
| Larval length (mm) | 11.6 | $3.37 \times 10^{-6}$ | 12.99 | $7.227 \times 10^{-6}$ | 23.046 | $3.951 \times 10^{-5}$ |
| Cocoon transverse axis length (mm) | 1.833 | 0.15 | 1.174 | 0.333 | 9.0135 | 0.029 |
| Fecundity (No.) | 6.524 | $6 \times 10^{-4}$ | 9.3175 | $1.222 \times 10^{-4}$ | 17.598 | $5 \times 10^{-4}$ |
| Pupae weight (g) | 1.96 | 0.129 | 2.174 | 0.108 | 5.61 | 0.132 |
| Cocoon longitudinal axis length (mm) | 13.86 | 0.055 | 4.883 | 0.006 | 9.965 | 0.019 |
| Cocoon weight (g) | 1.774 | 0.161 | 1.849 | 0.156 | 4.829 | 0.185 |

Principal component analysis (PCA) was performed for evaluating the global differences between the local *B. mori* breeds based on all the measured parameters. We used the PCA function from the *FactoMineR* package in two scenarios, firstly taking input from the entire data set (n=20 values/parameter/breed) and secondly taking input from the mean of each parameter (n=1 value/parameter/breed). Using the raw dataset, the results of PCA revealed relatively low explained variance between the breeds, with principal component 1 (PC1) of 29.5% and PC2 of 12.3%. Nonetheless, plotting these PCA results showed a clear separation between breeds mostly along the PC1 axis, with three distinct groups, one for the B1 breed, one for the JH3 breed, and one composed of the overlapping GB and ACH breeds (Figure 1A). Additionally, we used the *KODAMA* R package which implements an alternative algorithm for the PCA on large datasets. Interestingly, KODAMA results revealed that variance in the parameters dataset was best explained by fitting a k-means clustering model with two clusters. The results were visualized using the t-distributed stochastic neighbor embedding (t-SNE) method (Figure 1B), showing slight overlapping between the four breeds of *B. mori* and suggesting that the two clusters consist of B1-JH3 and GB-ACH. These results are similar to the previous PCA with regard to group proximity, but the KODAMA analysis indicates a closer relation between the B1 and GB breeds, as well as slightly greater variability for all breeds along both PC1 and PC2 axis.

**Figure 1.**
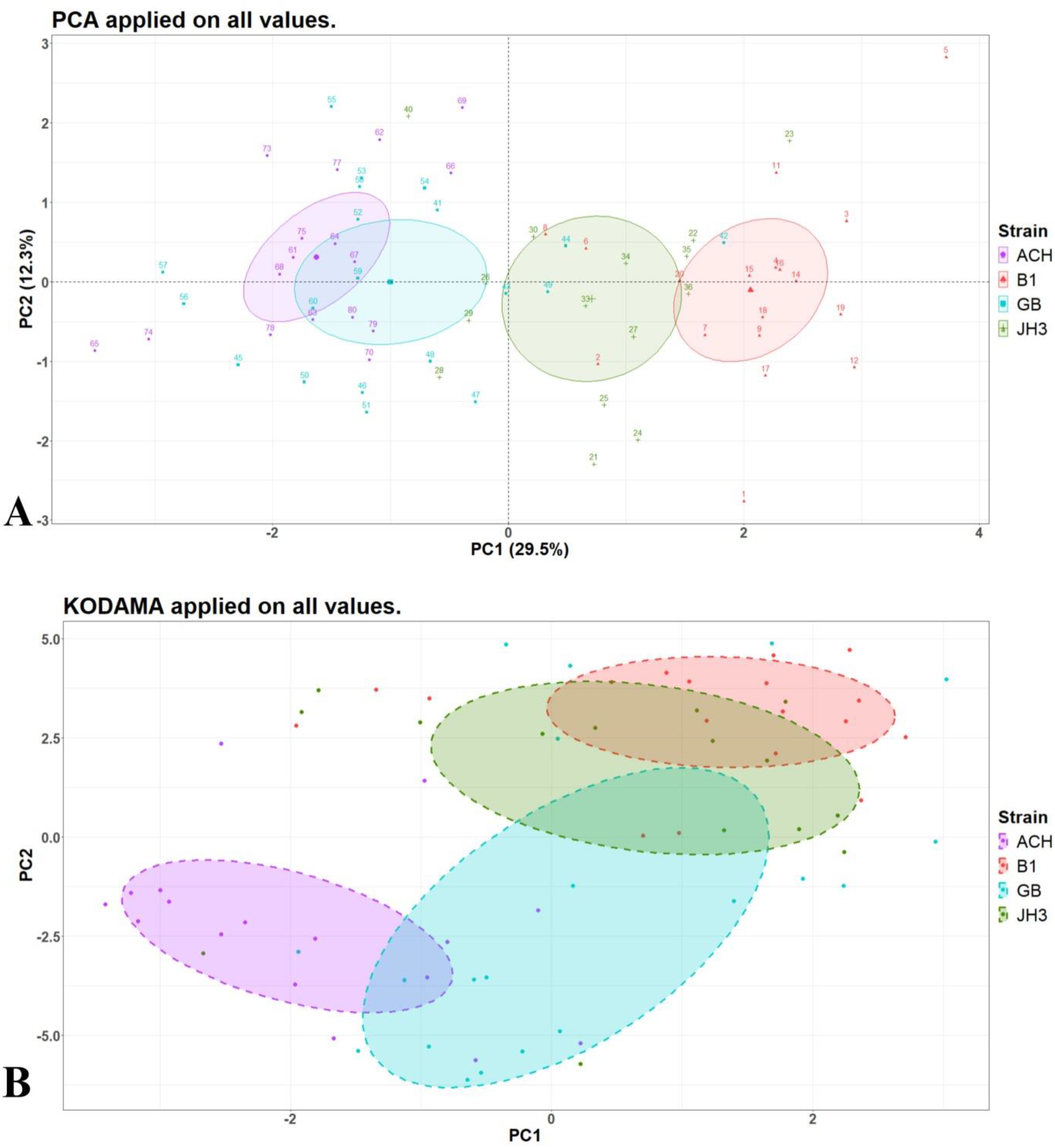
Graphical representations of PCA and KODAMA comparative analyses for all the measured parameters of *B. mori* development of the four local breeds. **A** – PCA of the entire dataset, plot realized with *fviz_pca_ind* function from the *factoextra* R package; **B** – KODAMA analysis of the entire dataset based on k-means clustering, plot realized with *method=“t-SNE”* argument in *KODAMA*.*visualization* function from the *KODAMA* R package.

Analysis of variance contributors revealed five measurements that explain the majority of variance between the breeds, these being correlated with the incubation period, larva characteristics (weight, length, and developmental instars), and the weight of the silk gland (Figure 2). These major contributors to variance (over 10% each), together with the “Fecundity” and “Cocoon longitudinal axis length” parameters were found to be significantly different in the ANOVA and Kruskal-Wallis tests. “Cocoon transverse axis length” was found significantly different only with the Kruskal-Wallis test.

**Figure 2.**
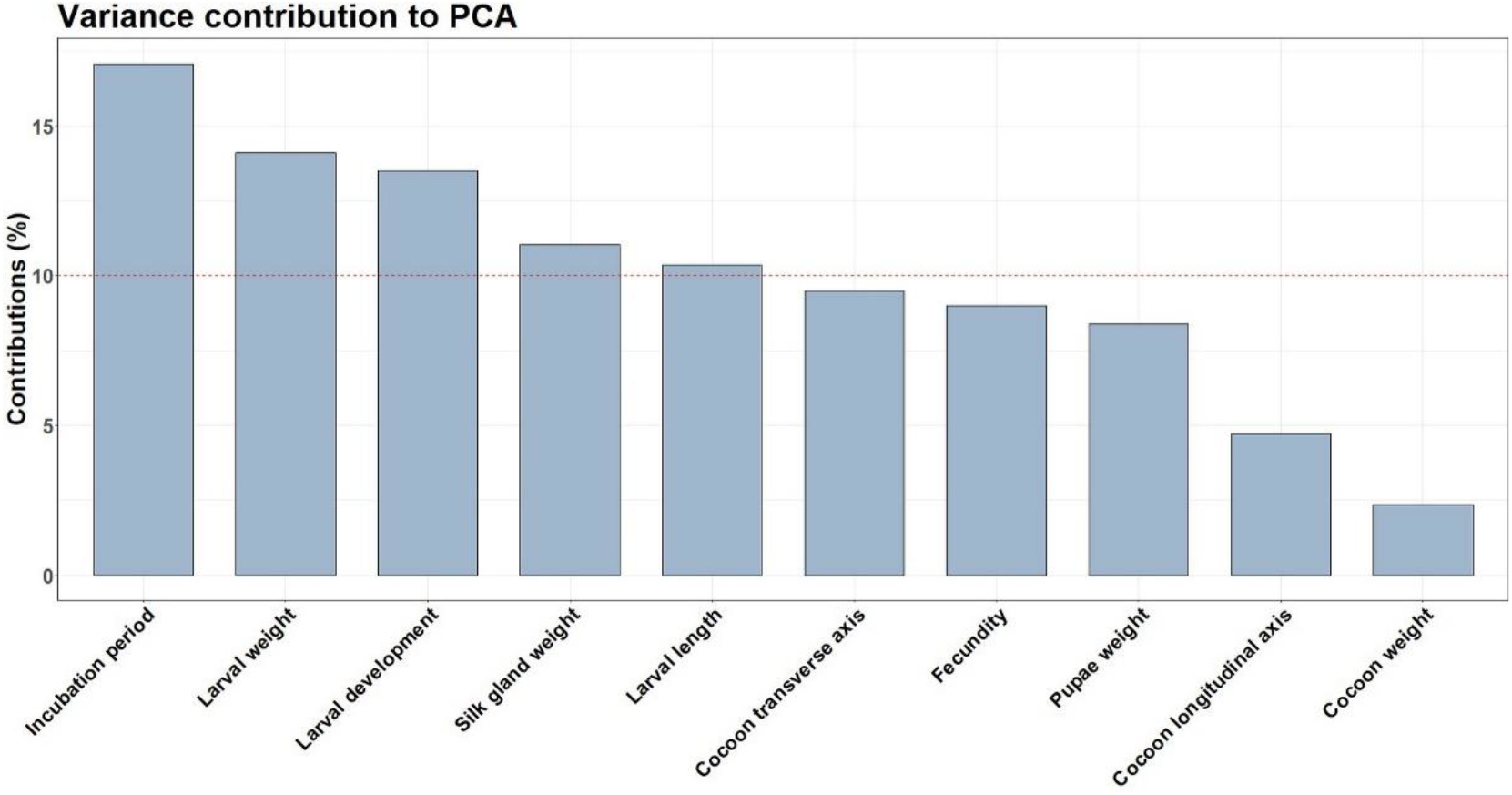
Barplot of parameters’ variance contributors within the entire dataset, realized with *fviz_contrib* function from the *factoextra* R package.

PCA of the parameters means dataset revealed a similar result to the entire dataset’s PCA, including group distribution along the PC1 axis and variance contributors (Figure 3). Rather interestingly, when using only the means of the measured parameters, 87% of the variance between the breeds was explained by PC1 (71.9%) and PC2 (15.9%).

**Figure 3.**
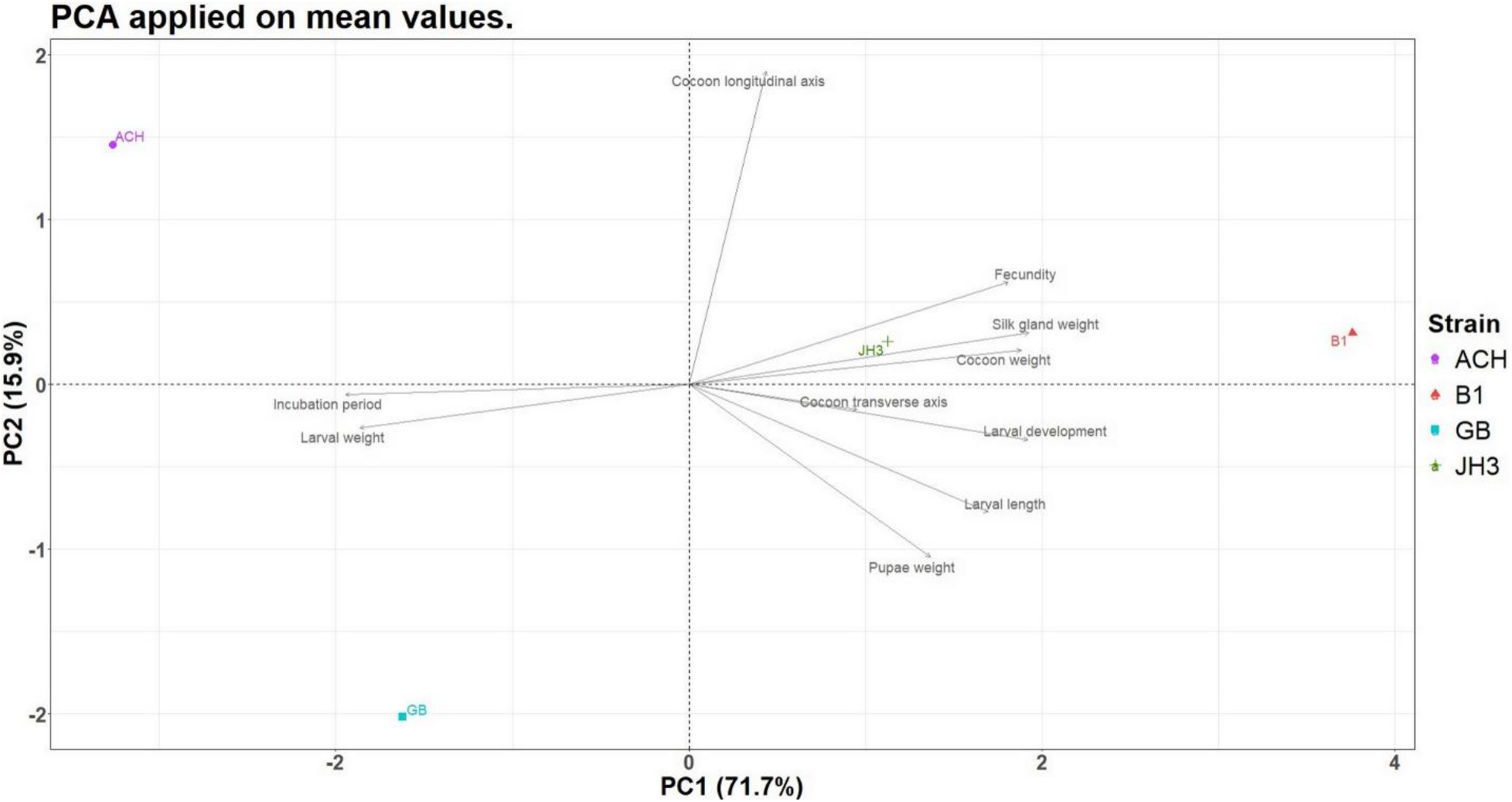
Biplot of individuals and variables corresponding to a dataset of measured parameters means, realized with *fviz_pca_biplot* function from the *factoextra* R package.

In order to exhaustively examine the ANOVA and PCA results, we performed a series of Tukey’s range (Tukey HSD) tests between the four *B. mori* breeds for each measured parameter. The statistical significance (p values) of each result is shown in Table 4 and graphically represented in Figure 4.

**Table 4.** Results (p values) of Tukey HSD tests between the four *B. mori* breeds for each measured.

| Statistical test | Tukey HSD test |  |  |  |  |  |
| --- | --- | --- | --- | --- | --- | --- |
| Comparison | B1 vs JH3 | B1 vs GB | B1 vs ACH | JH3 vs GB | JH3 vs ACH | GB vs ACH |
| Incubation period (days) | $3 \times 10^{-4}$ | $< 1 \times 10^{-7}$ | $< 1 \times 10^{-7}$ | $< 1 \times 10^{-7}$ | $< 1 \times 10^{-7}$ | 0.014 |
| Larval weight (g) | 0.127 | $< 1 \times 10^{-7}$ | $< 1 \times 10^{-7}$ | 0.073 | 0.046 | 0.99 |
| Larval duration (days) | 0.918 | 0.003 | $5 \times 10^{-4}$ | 0.032 | 0.007 | 0.899 |
| Silk gland weight (g) | 0.778 | 0.0035 | $9 \times 10^{-4}$ | 0.08 | 0.026 | 0.938 |
| Larval length (mm) | 0.102 | 0.0008 | $< 1 \times 10^{-7}$ | 0.47 | 0.014 | 0.263 |
| Cocoon transverse axis length (mm) | 0.986 | 0.999 | 0.283 | 0.988 | 0.179 | 0.253 |
| Fecundity (No.) | 0.925 | 0.002 | 0.999 | 0.028 | 0.873 | 0.002 |
| Pupae weight (g) | 0.571 | 0.618 | 0.084 | 0.998 | 0.731 | 0.579 |
| Cocoon longitudinal axis length (mm) | 0.374 | 0.054 | 0.118 | 0.845 | 0.944 | 0.995 |
| Cocoon weight (g) | 0.143 | 0.465 | 0.336 | 0.842 | 0.953 | 0.991 |

**Figure 4.**
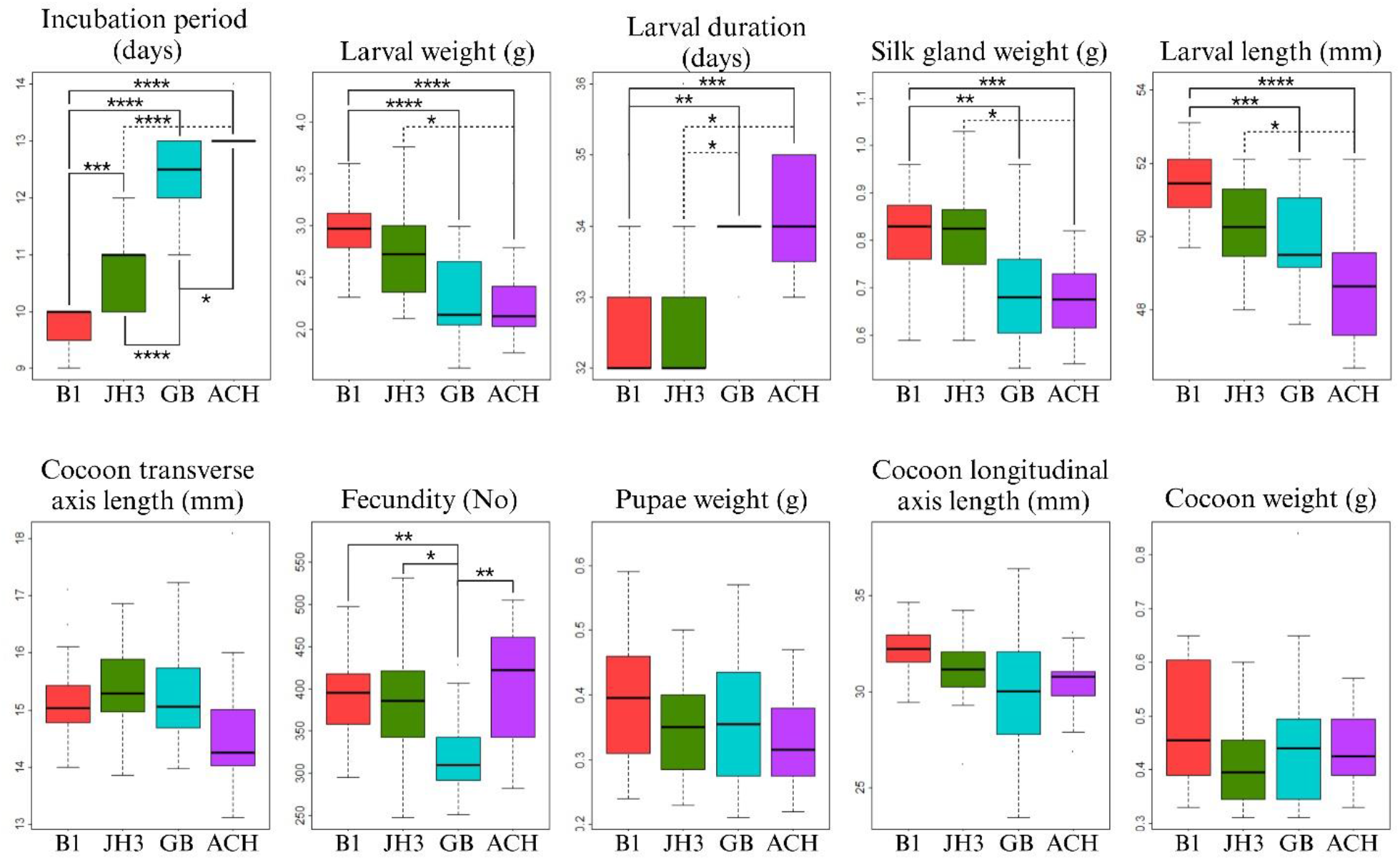
Boxplots of the measured traits corresponding to the four *B. mori* breeds. Statistical significance is shown as p value calculated based on the Tukey HSD test: * ≤ 0.05, ** ≤ 0.01, *** ≤ 0.001 and **** ≤ 0.0001. B1, JH3, GB and ACH breeds are represented by red, green, blue and purple boxplots, respectively.

The ANOVA analysis and variance contributors of PCA are reflected in the Tukey HSD tests results, as the most significantly different parameters across the breeds, and the major contributors to variance show statistical significance in multiple comparisons. One notable example is the “Incubation period” parameter, where all four breeds were found to be statistically different. Moreover, visual analysis of the boxplots supports the results of the PCA and KODAMA analyses, with an observable separation between the B1-JH3 and GB-ACH breeds (Figure 1 and Figure 3). This grouping of breeds is especially visible for the four largest contributors to variance, namely the “Incubation period”, “Larval weight”, “Larval duration”, “Silk gland weight”, and “Larval length” parameters.

The results of Tukey HSD test were validated with the Games-Howell nonparametric post-hoc test which does not assume equal variances between the variables. Overall, the profile of significant differences between the breeds was maintained following nonparametric testing. We observed three notable exceptions: lack of statistical significance between JH3 and GB breeds for “Larval duration” parameter (p = 0.032 for Tukey HSD test and p = 0.105 for Games-Howell test), statistical significance between B1 and JH3 breeds for “Larval length” parameter (p = 0.102 for Tukey HSD test and p = 0.037 for Games-Howell test) and statistical significance between B1 and ACH breeds for “Cocoon longitudinal axis length” parameter (p = 0.118 for Tukey HSD test and p = 0.008 for Games-Howell test). In contrast, the nonparametric testing showed that B1 breed is not different to GB breed regarding the “Cocoon longitudinal axis length”, with a p value of 0.116, compared to 0.054 from the Tukey HSD test. Regarding the “Fecundity”, although the Games-Howell test returned a p = 0.06 when comparing JH3 and GB breeds, we consider that the difference is significant since Tukey HSD test returned a p = 0.028.

## 4. Discussions

The larvae of *B. mori* exhibit a great capacity to exploit the mulberry leaf proteins in order to produce silk (Saviane et al., 2014). Considering the scientific and economic value of various *B. mori* breeds, it is of strategic importance to evaluate and optimize both the production parameters and essential biological features of the silkworm developmental stage. The exhaustive traits examination of the four *B. mori* breeds is critical for optimizing silk production and obtaining sustainable sericulture practices, but also for paving the future of biotechnology, material, or pharmaceutical sciences.

The biological and economic parameters of the *B. mori* larvae and adults that were fed with an artificial diet were assessed in the November-December period. Yin et al. (2023) argued that feeding the silkworms with an artificial diet negatively impacts some of their traits, therefore affecting their economic value. Even by supplementing the artificial diet with bee pollen, the measured parameters were not significantly improved (Moise et al., 2020). On the other hand, adding Zinc to the artificial diet led to significant differences in terms of silk glands and cocoon weight, but also related to the cocoon shell (Benţea et al., 2012). In the present study, we explored various parameters of the target breeds, i.e., the incubation period, larval weight, larval duration, silk gland weight, larval length, cocoon transverse axis length, fecundity, pupae weight, cocoon longitudinal axis length, and cocoon weight. Since the measured parameters had normal distribution, for data comparisons we employed specific parametric statistical tests, more specifically one-way ANOVA and Welch’s ANOVA test; the Kruskal-Wallis nonparametric test was also employed as a less restrictive analysis method and because of the relatively small number of individuals selected for each breed. Among the measured traits, “pupae weight” and “cocoon weight” did not reach statistical significance. The “cocoon transversal axis length” was evaluated as significantly different only with the nonparametric test. The results referring to cocoons could indicate that there is a sizable interindividual variability of these parameters within certain breeds, possibly because of biased individual selection or the particular artificial diet.

The traits comparison between *B. mori* breeds represents the base of any breeding program. In order to have a snapshot of the global differences noticed for the four breeds, PCA, KODAMA and Tukey HSD tests were performed, using all measured data or their means. PCA analyses have been intensively used for interpreting data drawn from genome-wide association studies (GWAS) which consist of large datasets of both continuous and categorical variables. Although less frequent, PCA has been successfully implemented for investigating possible similarities between various groups based on the observed phenotypes. For example, PCA was used in research to correlate host-commensal microbiota interactions in *D. melanogaster* laboratory breeds (Han et al., 2017), or, in more clinical-orientated studies, to assess possible relations between multiple clinical phenotypes of patients suffering from chronic obstructive pulmonary disease (Roy et al., 2009; Burgel et al., 2010), obstructive sleep apnea (Vavougios et al., 2016) or Alzheimer disease (Zeitzer et al., 2013).

Our PCA allowed along the PC1 axis a noticeable distinction between B1, JH3 and an apparent group formed by GB and ACH, which was confirmed using both all and mean values corresponding to the measurements of the biological parameters. Using the mean values, the percent of the variance between the breeds explained by PC1 raised to 87% from a mere 29.5% calculated with complete data. This result might suggest a possible benefit when sample means are used, as large standard deviations in the populations result in overlapping of distribution tails, as can be observed from Figure 4 and the corresponding histograms (Supplementary Figure 1). In a study arguing the importance of breed selection for the success of any further enterprises, Bhat et al. (2018) compared 18 silkworm breeds and found significant differences among the traits with economical relevance (Bhat et al., 2018). In our hands, KODAMA analysis also revealed the likely distinct group formed by GB and ACH. Since both these breeds are producing naturally colored silks, a feature that comes with particular biochemical and genomic characteristics (Ma et al., 2016), it is tempting to ascribe to this particularity their relative similarity concerning some biological parameters. As for B1 and JH3, although nowadays the former breed can be considered as Romanian, both share a common Japanese origin (Furdui et al., 2014).

Analysis of variance contributors to PCA revealed that five parameters, namely “Incubation period”, “Larval weight”, “Larval duration”, “Silk gland weight” and “Larval length” explain more than 10% of differences between the four *B. mori* breeds studied. Similar investigations on biological traits were performed on various *B. mori* breeds by independent groups. Zanatta et al. (2009) found noticeable differences between 16 Chinese or Japanese breeds regarding cocoon length, cocoon weight, raw silk percent and larval body weight (Zanatta et al., 2009). In addition, Bhat et al. (2018) identified variability between 18 silkworm breeds for phenotypic traits such as fecundity, hatching percentage, survival percentage, single cocoon weight, single shell weight and silk filament length (Bhat et al., 2018). Results of this type suggest that certain biological traits are specific for particular *B. mori* breeds and selective breeding strategies can enhance desired traits. Such phenotype selections were demonstrated by Zamani et al. (2019) when breeding selection between Japanese and Chinese breeds resulted in improved “Cocoon shell percentage” and “Cocoon shell weight” (Zamani et al., 2019).

Environmental factors such as temperature and humidity have been shown to influence various phenotypes of *B. mori* breeds (Hussain et al., 2011; Rahmathulla and Suresh, 2012). Modifications in temperature and humidity resulted in significant differences in hatchability, pupation and larval survival of multiple inbred *B. mori* lines (Hussain et al., 2011). The authors of this study identified that a temperature of 25-26°C and a humidity of 70-80% is optimal for silkworm husbandry (Hassain et al., 2011). Moreover, nutrients in food, such as vitamin C, vitamin E, and vitamins from the B complex, have been shown to influence key biological traits of *B. mori* individuals (Kanafi et al., 2007). Taken together, these studies show evidence of biological traits enhanced by environment modifications, which, paired with our findings about the genotypes of four *B. mori* breeds, might result in even further enhancing specific traits of interest. For example, to minimize the incubation period in our breeds, a combined strategy of selective breeding and optimization of environmental factors might be recommended.

Moreover, a complex review of the environmental impact on silkworm phenotypes discusses how temperature and humidity on growth and development (Rahmathulla and Suresh, 2012). Interestingly, the authors identified that the environmental contribution is higher for pupation percentage, cocoon weight parameters and shell ratio than the genetic contribution (Rahmathulla and Suresh, 2012). Additionally, raw silk percentage is influenced equally by the two types of factors, and filament length is influenced especially by genetic factors (Rahmathulla and Suresh, 2012). The variance contribution to PCA found in our study revealed that the “Cocoon weight” parameter explains the least variability between the four *B. mori* breeds, and this observation is consistent with the findings of Rahmathulla and Suresh (2012), as this biological trait is little affected by the genotype. In contrast, the “Cocoon weight” parameter has been identified to be highly correlated with shell weight (R=0.961), larval length (R=0.945), shell ratio (R=0.861) and larval duration (R=0.648) phenotypes (Pradeep et al., 2007). In our analysis, the “Larval weight” and “Larval duration” parameters were the second and respectively third highest contributors to variance, but the “Cocoon weight” parameter explained very little differences between the *B. mori* local breeds.

## 5. Conclusion

This study implies a comprehensive effort to characterize four *B. mori* breeds originating from Romania (B1, GB), Japan (JH3), and China (ACH) in terms of their economic relevant traits. The exhaustive characterization of certain parameters of *B. mori* breeds promotes the development of the target breeding programs. To register significant progress in gathering knowledge and utilization of *B. mori* in various fields, interdisciplinary collaboration is required.

## Acknowledgments

This work was supported by the Ministry of Agriculture and Rural Development of Romania through project ADER 24.1.1/2023.

