## Supplementary Figures for "Novel insights into the economic relevant phenotypes of *Bombyx mori* strains from Romanian breeding centres"

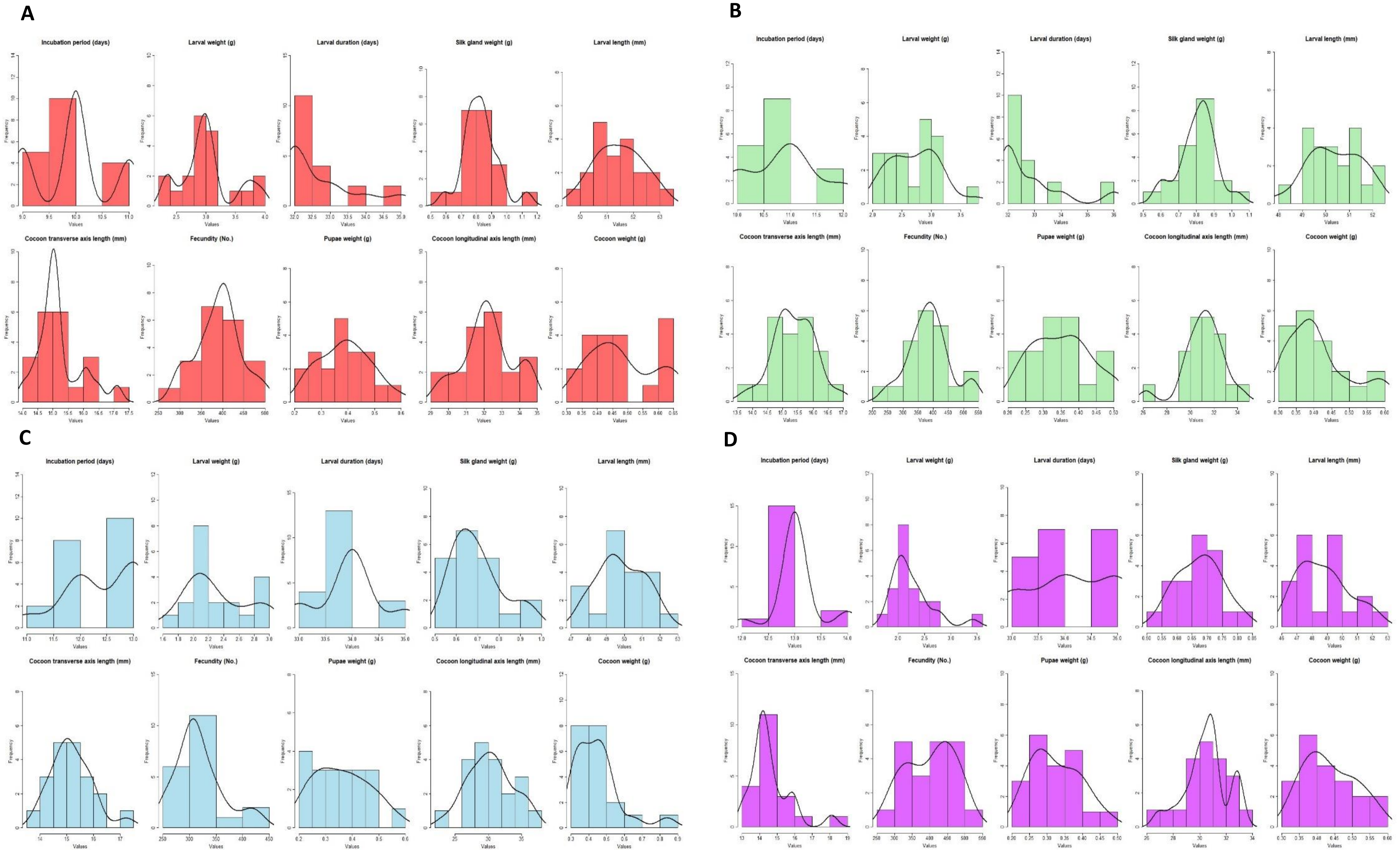


**Supplementary Figure 1.** Histograms of each measured biological parameters corresponding to the four local *B. mori* strains of interest: A-B1, B-JH3, C-GB, D-ACH. Histograms display the frequency of raw data using custom bins for each parameter and densities are shown as a black line for better graphical representation. All graphs were generated in RStudio.

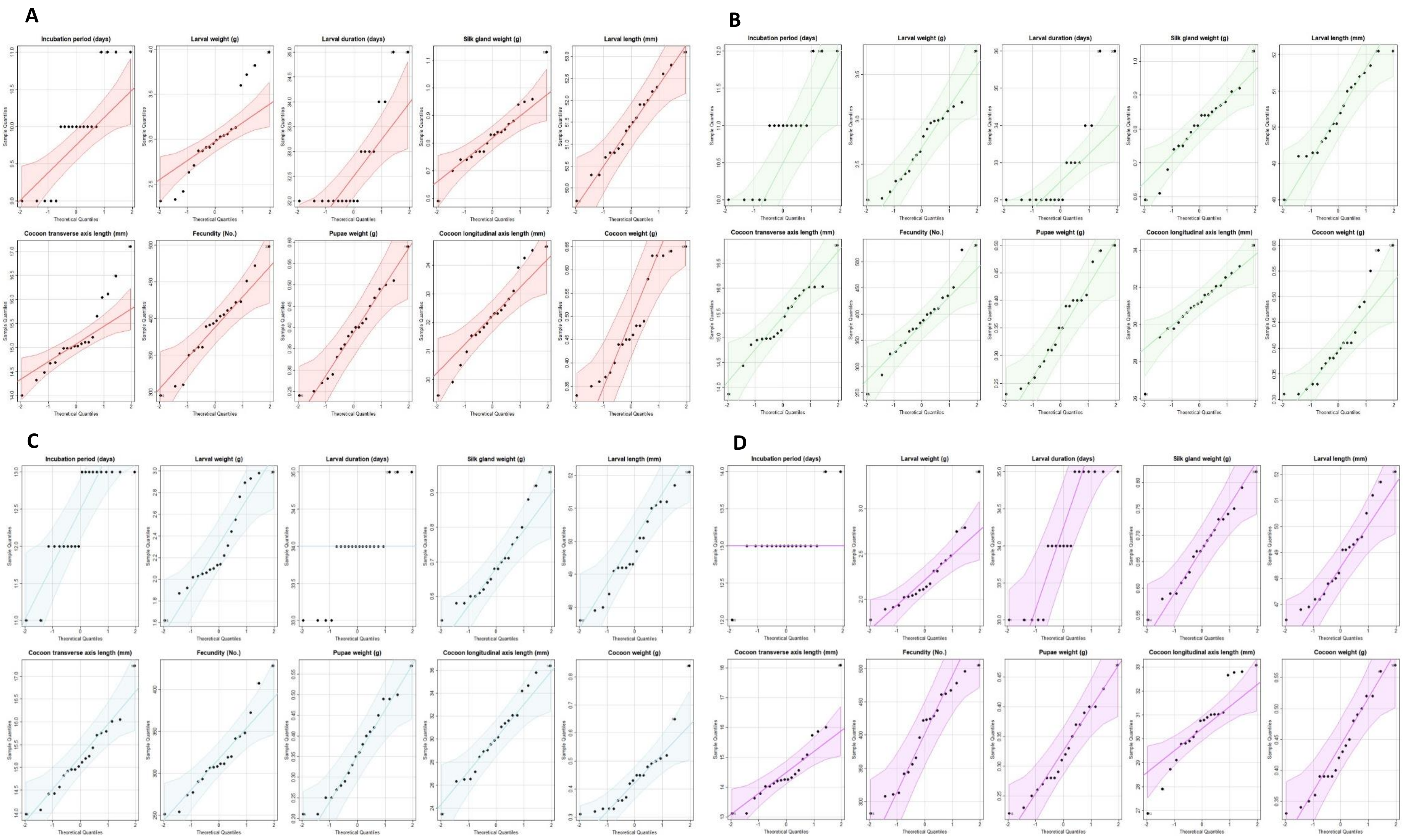


**Supplementary Figure 2.** Quantile-quantile (Q-Q) plots of each measured biological parameters corresponding to the four local *B. mori* strains of interest: A-B1, B-JH3, C-GB, D-ACH. Each graph displays the linear relation between quantiles drawn from the parameter distribution and the normal distribution. All graphs were generated using the *qqPlot()* function from the *EnvStats* R package in RStudio.
